# Morphological characteristics of the forelimb flexor digitorum muscles and their distinctive innervation pattern in koalas (*Phascolarctos cinereus*): implications for function and morphogenesis

**DOI:** 10.64898/2026.09.05.749653

**Authors:** Daiki Shigemitsu, Saori Anetai, Hidaka Anetai, Jaliya Kumaratilake, Chris Leigh, Kounosuke Tokita, Sayaka Tojima

**Author notes:** **Correspondence to:** Dr. Saori Anetai Department of Physical Therapy, Faculty of Health and Medical Care, Saitama Medical University, 981 Kawakado, Moroyama-machi, Iruma-gun, Saitama 350-0496, Japan.

## Abstract

Koalas (*Phascolarctos cinereus*) are marsupials highly specialized for arboreal life and are considered a representative example of convergent evolution with primates, which are also adapted to arboreal habitats. The forelimb digital flexors play a crucial role in arboreal locomotion by flexing the digits and enabling the claws or fingers to engage with tree trunks and branches. Although the morphology and function of the digital flexors have been extensively examined in primates, anatomical information on koala digital flexors has not been sufficiently updated in modern anatomical studies. In this study, we examined the digital flexor muscles and their innervation in four forelimbs from three koalas to clarify both shared and distinctive structural features among arboreal mammals. The flexor digitorum profundus (FDP) was markedly developed, formed a U-shaped muscle belly, and received dual innervation from the median and ulnar nerves. In contrast, the flexor digitorum superficialis (FDS) was small, arose from the superficial surface of the FDP, was partially enclosed by the FDP muscle belly, and gave rise to extremely slender tendons inserting into digits II–V. The branch supplying the FDS arose from the main trunk of the median nerve near the carpal region, ascended proximally in a recurrent course, and terminated within the FDS. This unusual course may be consistent with a morphogenetic process underlying FDS formation. These findings indicate that the koala digital flexors are characterized by a markedly developed FDP, a relatively reduced FDS, and a distinctive innervation pattern of the FDS. Thus, even among mammals adapted to arboreal life, koalas and primates appear to have achieved arboreal adaptation through different morphological specializations within the forearm digital flexor system.

## 1. Introduction

Marsupials are mammals that inhabit mainly Australia and South America and are characterized by nurturing their underdeveloped young in a pouch located on the mother’s abdomen (Kemp, 2004). They exhibit diverse locomotor modes associated with various habitats, such as grasslands and woodlands. For example, kangaroos and wombats are both ground-dwelling marsupials: kangaroos move by hopping on their hind limbs at higher speed or by pentapedal locomotion at lower speed, whereas wombats walk quadrupedally (Cartmill et al., 2019; O’Connor et al., 2014). By contrast, koalas (*Phascolarctos cinereus*) are primarily arboreal and feed mainly on eucalyptus leaves. Koalas are marsupials specialized for arboreal life and possess distinctive morphological characteristics associated with this lifestyle, likely reflecting the functional demands of arboreal living.

We recently examined the gluteal and femoral flexor muscles in koalas and discussed their morphology from the perspective of functional demands associated with climbing and clinging to tree trunks. We revealed a high degree of independence between the two-layered gluteus medius in koalas and suggested that this feature may be related to enhanced hip-joint extension and abduction functions, thereby enabling koalas to ascend, descend, and cling to tree trunks of various thicknesses (Tojima et al., 2022). Focusing on the forearm muscles, the digital flexors are likely to play a crucial role in climbing and clinging to tree trunks because they enable the digits to flex and the claws to engage with the trunk. Therefore, the digital flexors are considered essential for arboreal locomotion, implying a close functional anatomical relationship between the morphology of the digital flexors and claw function in koalas.

Despite their likely functional importance, anatomical studies on marsupial muscles have been limited. Macalister (1870, 1872) provided the earliest descriptions of the muscular anatomy of marsupials, including koalas, wombats, and Tasmanian devils; however, these early works were brief and consisted mainly of lists of muscular features. Young (1882), building on Macalister’s observations, provided an updated and more detailed description based on dissections of three koala specimens, although no illustrations were included.

Later, Sonntag (1922) conducted myological comparisons among the wombats, koalas, and phalangers from a taxonomic perspective and was the first to include some illustrations; however, this study did not address functional morphological considerations. Thus, existing studies on koala muscle morphology remain limited, and modern investigators must rely largely on outdated and insufficient anatomical descriptions. Lee and Carrick (1989) noted that caution is warranted when comparing Sonntag’s descriptions of koala limb musculature with modern anatomical accounts because of discrepancies in terminology. Richards et al. (2023) further pointed out that interpretations and comparisons based on these early myological descriptions are complicated by outdated anatomical terminology and historical constraints on published illustrations. Therefore, the muscular anatomy of koalas requires re-examination by modern morphologists, potentially allowing us to consider them from viewpoints of functional anatomy and adaptation.

Among mammals, primates represent an example of morphological adaptation to arboreal locomotion. Research on the morphology and function of the digital flexors has been extensively conducted in primates, which are also highly arboreal, and the morphological characteristics of the muscles involved in tree climbing and branch grasping have been elucidated in detail (Diogo and Wood, 2011; Emura et al., 2020, 2023; Leischner et al., 2018; Youlatos, 2000). From an evolutionary perspective, koalas and primates are considered a representative case of convergent evolution because both are adapted to arboreal life. Convergent evolution is the evolutionary process by which organisms from different lineages acquire similar morphological and functional characteristics as they adapt to similar environments and lifestyles (Romer, 1997). The digital flexors, which are essential for arboreal locomotion, may exhibit morphological similarities in koalas and primates.

Therefore, a comparison of the morphological characteristics of the digital flexors in koalas and primates may help clarify morphological adaptations to arboreal locomotion. In this study, we conducted a detailed examination of the digital flexors in koalas, an arboreal marsupial, and identified both shared and unique structural characteristics relative to primates for functional anatomical and adaptative considerations.

## 2. Materials and Methods

Four forelimbs from three koalas were examined in this study (Table 1). Three forelimbs from two specimens were obtained from the University of Adelaide, and one forelimb was from the collection of Tokyo Ariake Medical University. Although the ages of all three koalas were unknown, one specimen was presumed to be adult (Ph-1) and the other two were presumed to be juveniles on the basis of body size. Before dissection, the specimens were fixed in formalin and subsequently preserved in 50% ethanol with glycerin. This study was conducted in accordance with the Guiding Principles for the Care and Use of Animals of the American Association of Anatomists. No animals were euthanized specifically for this study.

**Table 1.** Specimens examined in the present study.

| Genera/Species | Specimen no. | Side | Body size (mm) | Remark |
| --- | --- | --- | --- | --- |
| *Phascolarctos cinereus* | U-Adelaid-1 | Left | 820 | U-Adelaid-1 |
|  | U-Adelaid-2 | Right |  |  |
|  | Ariake-1 | Left | 585 | Ariake-1 |
|  | Ariake-2 | Left | 560 | Ariake-2 |

In the present study, we focused on the forelimb digital flexors, specifically the flexor digitorum superficialis (FDS) and flexor digitorum profundus (FDP). Additionally, the branches from the median and ulnar nerves that innervated these muscles were also examined in detail because innervation can provide important insights into considering muscle homology and morphogenesis (Anetai et al., 2026; Emura et al., 2020, 2023; Koizumi, 2019, 2022, 2023; Sakuraya *et al*., 2023; Velez-Garcia et al., 2026). The koalas were dissected according to a standard procedure for human dissection in Japan (Yamada & Mannen, 1985). Muscle identification and nomenclature were based on the *Nomina Anatomica Veterinaria* (6th ed.). First, the skin and subcutaneous tissues were removed from the upper arm to the digits to expose the forearm muscles including their insertion tendons. The morphological characteristics of the superficial forearm flexors, including the flexor carpi radialis, palmaris longus, and flexor carpi ulnaris, were then documented using photographs and detailed drawings. Thereafter, the palmaris longus and palmar aponeurosis were detached and reflected to allow detailed observation of the morphology of the FDS and FDP as well as their innervation. Subsequently, the forearm flexor group was detached en bloc from its origin to examine the attachment sites. The origin, course, and distribution of the innervating branches were examined using an operating microscope as necessary.

## 3. Results

In all examined forelimbs, the pronator teres, flexor carpi radialis, palmaris longus, flexor carpi ulnaris, FDS, and FDP were identified (Table 2), and their innervation were characterized in detail. The morphological characteristics of these muscles were consistent across the specimens examined, with no apparent differences associated with body size, species, or sex. The following sections describe the forelimb digital flexors from two perspectives: their detailed muscle structure and innervation.

**Table 2.** Morphological characteristics of the muscles examined.

| Muscle | Origin | Insertion | Innervation |
| --- | --- | --- | --- |
| Anconeus | Humerus epicondylus medialis | Olecranon | Ulnar nerve |
| Flexor carpi radialis | Humerus epicondylus medialis | Base of the 2nd metacarpal bone | Median nerve |
| Flexor carpi ulnaris | Humerus epicondylus medialis | Pisiform | Ulnar nerve |
| Flexor digitorum superficialis | Surface of distal 1/2 flexor digitorum profundus | Base of the 2nd to 5th middle phalanx | Median nerve |
| Flexor digitorum profundus | Humerus epicondylus medialis, | Base of the 1st to 5th distal phalanx | Median and Ulnar nerve |
|  | Proximal 1/2 of ulna, Middle 1/3 of radius |  |  |
| Palmaris longus | Humerus epicondylus medialis | Palmar aponeurosis | Median nerve |
| Pronator teres | Humerus epicondylus medialis | Middle 1/3 of radius | Median nerve |

### 3.1. Morphology of the forelimb digital flexors

#### 3.1.1. Flexor digitorum superficialis (FDS)

The FDS originated from the distal half of the superficial surface of the FDP and was partially enclosed by the FDP muscle belly (Figure 1). The FDS belly divided into four parts, each giving rise to a distal tendon that extended towards digits II–V. All four tendons were extremely thin. The FDS tendons were perforated by the FDP tendons and inserted into the middle phalanges of digits II–V (Figure 2). The muscle was innervated solely by the median nerve (see below for details).

**Figure 1.**
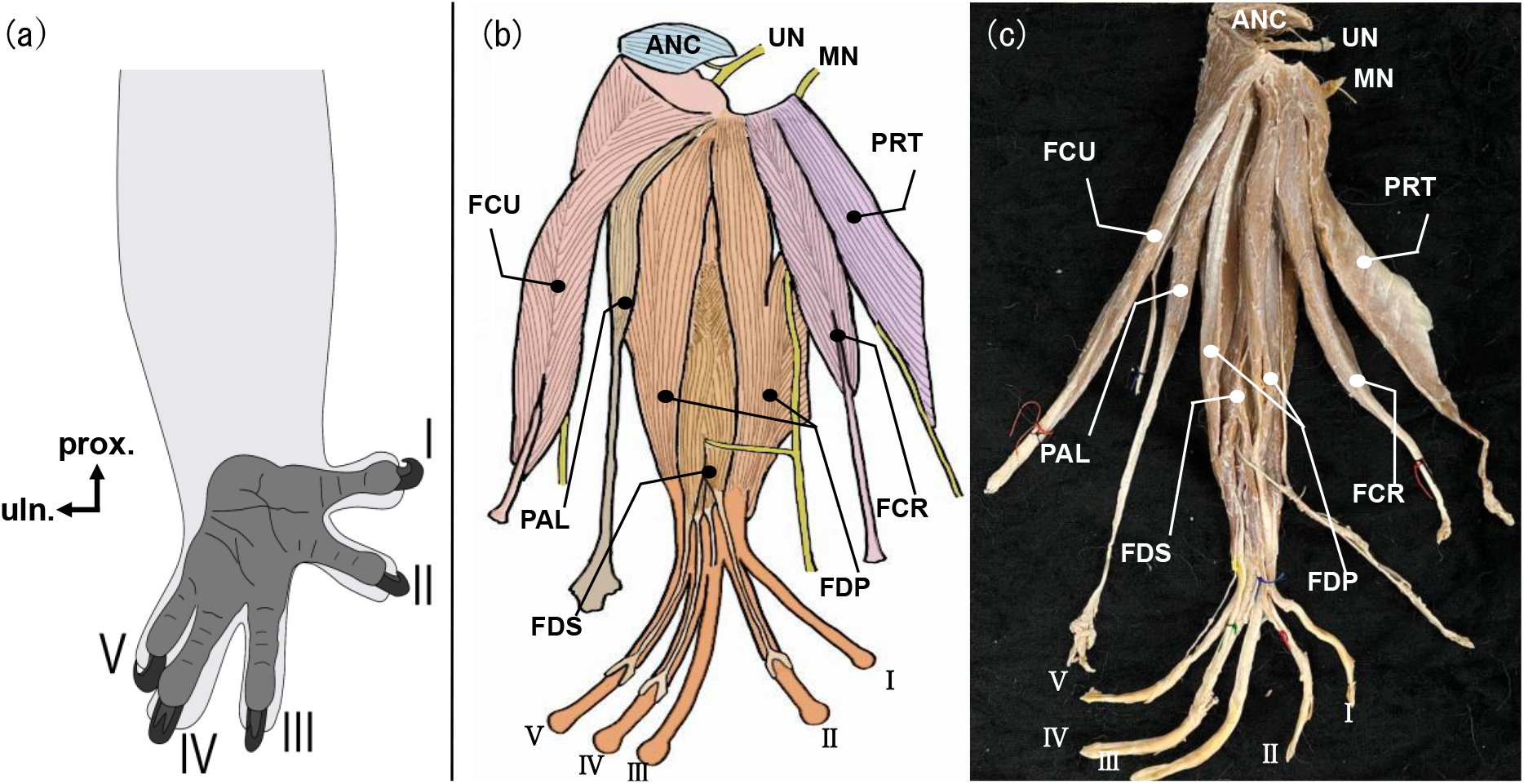
Morphology of the forearm flexor muscles (superficial aspect) The left forearm flexor muscles examined are presented. (a) Palmar view. Digits I and II are positioned opposite digits III–V as illustrated. (b) Accurate drawing of the forearm flexor muscles. (c) Photograph of the forearm flexor muscles. ANC, anconeus; FCR, flexor carpi radialis; FCU, flexor carpi ulnaris; FDS, flexor digitorum superficialis; FDP, flexor digitorum profundus; PAL, palmaris longus; PRT, pronator teres; MN, median nerve; UN, ulnar nerve.

**Figure 2.**
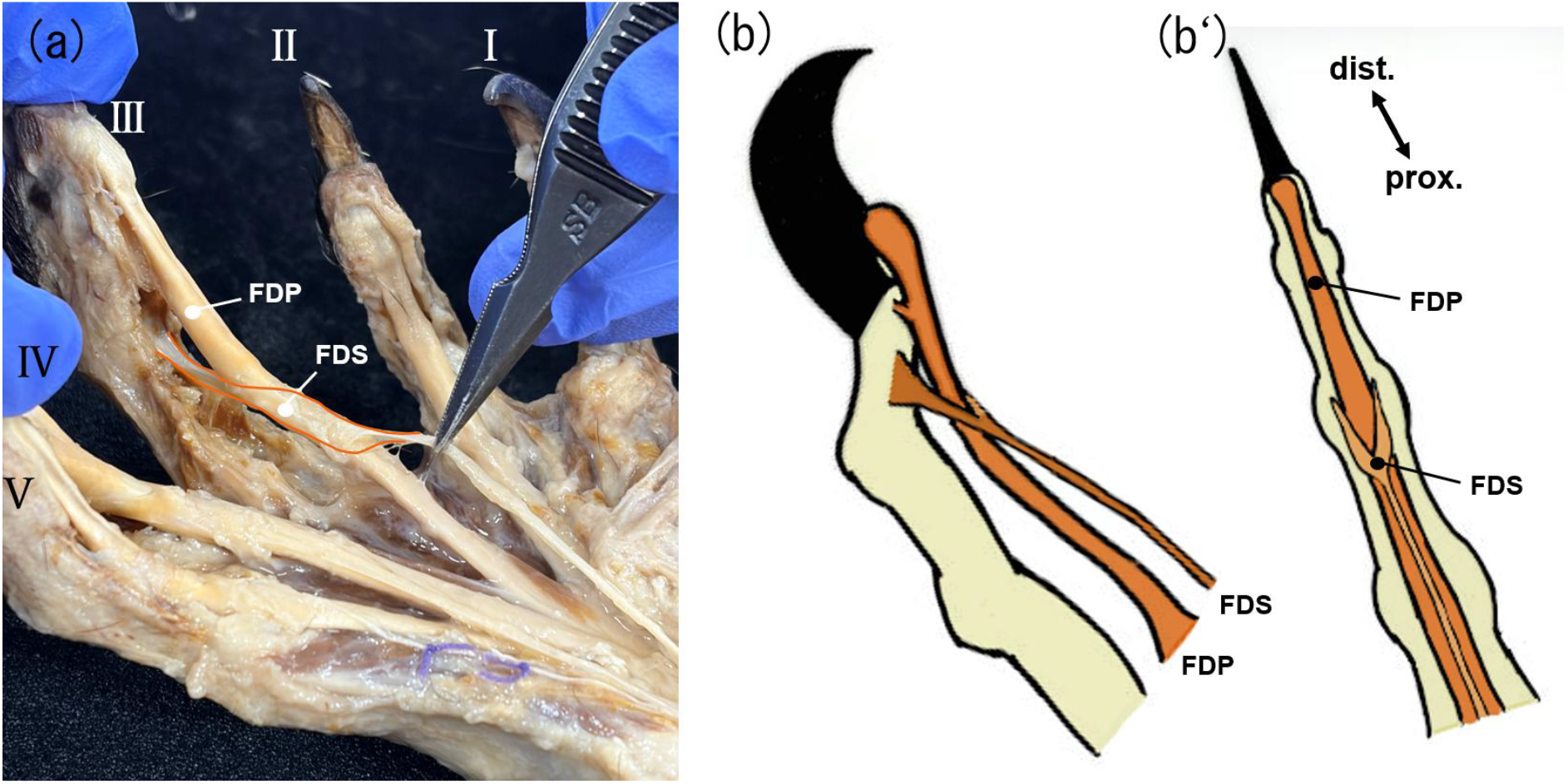
Morphology of the FDS and FDP tendons. The positional relationship between the FDS and FDP tendons is shown in a photograph (a) and schematic drawings (b) and (b’). (b) Lateral view. (b’) Palmar view. FDS, flexor digitorum superficialis; FDP, flexor digitorum profundus.

#### 3.1.2. Flexor digitorum profundus (FDP)

The FDP had the largest muscle belly among the forearm flexors. It originated from the medial epicondyle of the humerus, the proximal two-thirds of the ulna, and the middle one-third of the radius, and gave rise to two radial and three ulnar tendons (Figure 1). These tendons passed through the carpal tunnel, extended to digits I–V, penetrated the thin FDS tendons within the tendon sheath near the proximal phalanges, and inserted into the bases of the distal phalanges of digits I–V (Figure 2). The muscle belly of the FDP was folded into a U-shaped configuration, forming a concavity on its superficial surface. The FDS was located within and originated from this concavity. No independent flexor pollicis longus (FPL) belly was observed. The FDP was innervated by both the median and ulnar nerves (see below for details).

### 3.2. Course and branching pattern of the median and ulnar nerves

#### 3.2.1. Median nerve

The median nerve passed through the supracondylar foramen of the humerus to emerge in the flexor region, and first divided into two branches proximal to the forearm (Figure 3). One branch ran medially, gave off several muscular branches to the FDP, and finally communicated with a branch derived from the ulnar nerve. These muscular branches coursed along the superficial surface of the FDP and terminated to this muscle. The other branch, regarded as the main trunk, continued to the palm and gave off several branches along its course. From the proximal part of this main trunk, muscular branches were given off to the palmaris longus, pronator teres, flexor carpi radialis, and the superficial portion of the FDP. In contrast, the branch supplying the FDS diverged from the distal part of the main trunk near the carpal region and ascended proximally to the superficial surface of the FDS (Figures 3). This branch finally divided into several twigs that terminated in the respective muscle bellies for digits II–V (Figure 4).

**Figure 3.**
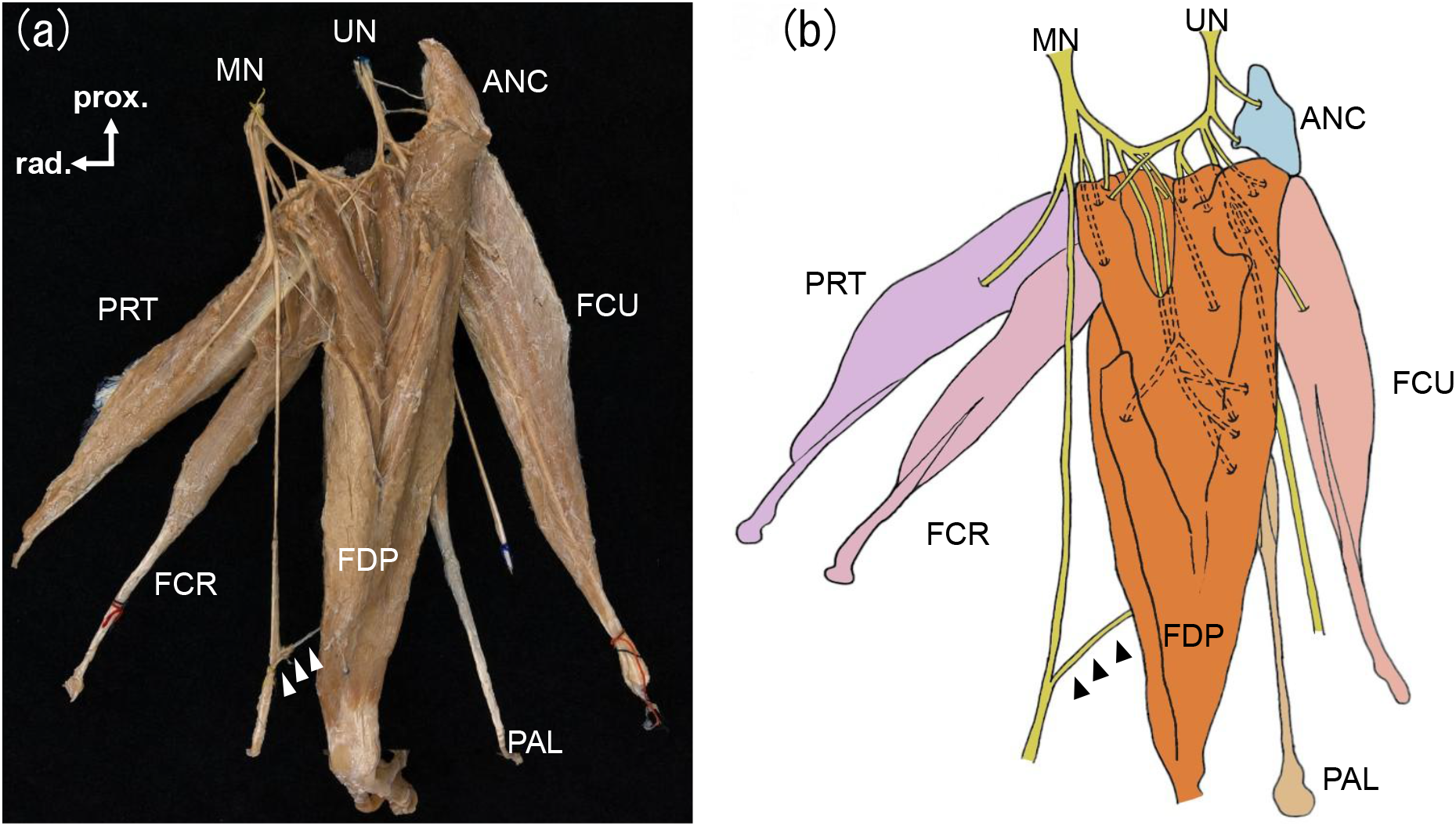
Innervation of the forearm flexor muscles (deep aspect) The courses and distributions of the median and ulnar nerves are shown on the deep aspect of the forearm flexor muscles. (a) Photographs of the innervating branches arising from the median and ulnar nerves and innervated muscles. (b) Accurate drawing. White and black arrowheads indicate the branch innervating the FDS. ANC, anconeus; FCR, flexor carpi radialis; FCU, flexor carpi ulnaris; FDS, flexor digitorum superficialis; FDP, flexor digitorum profundus; PAL, palmaris longus; PRT, pronator teres; MN, median nerve; UN, ulnar nerve.

**Figure 4.**
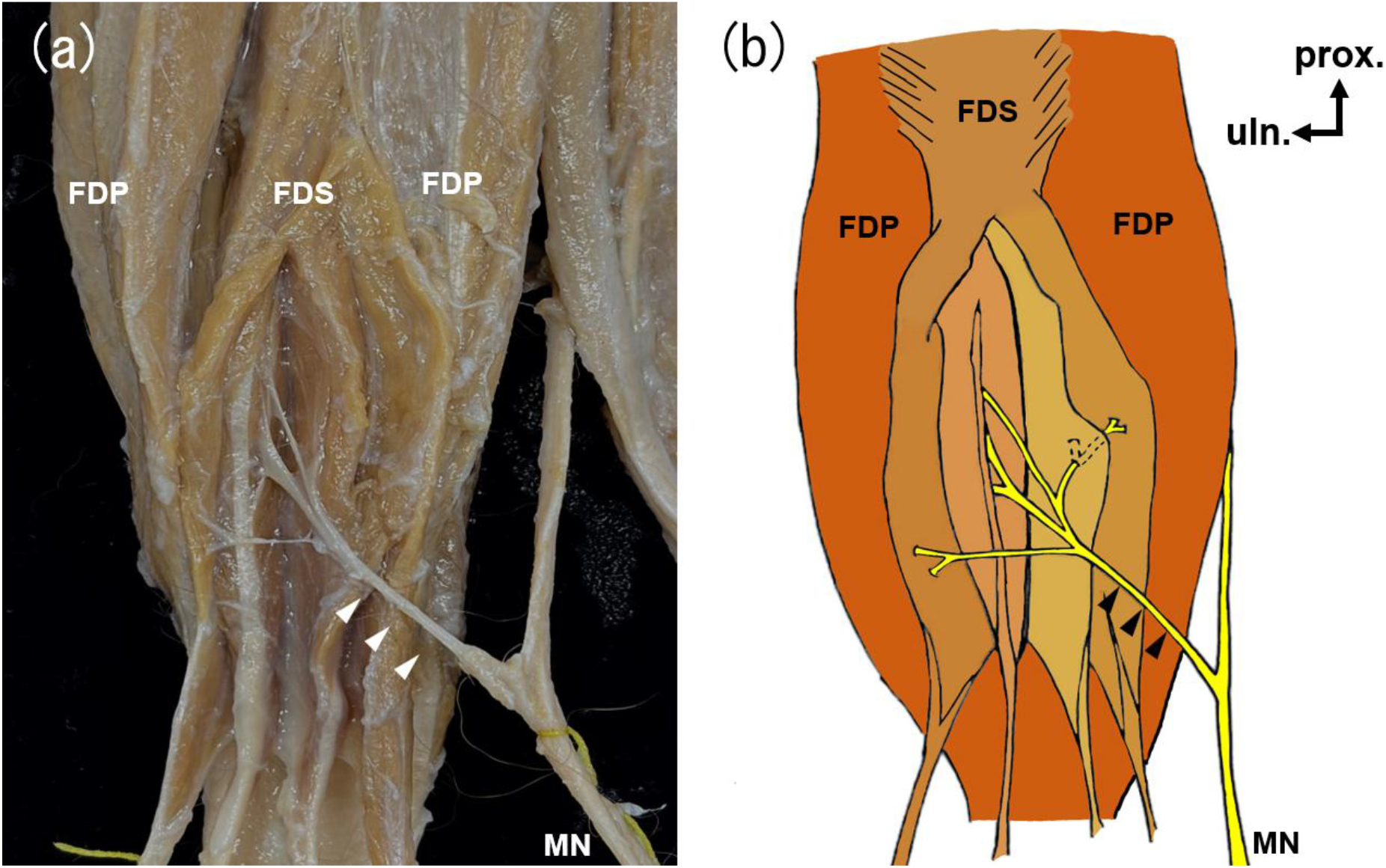
Detailed innervation pattern of the FDS (superficial aspect) The FDS muscle bellies to digits II and V were partially reflected to expose branches arising from the median nerve and entering the FDS bellies to digits II–V (a). (b) Accurate drawing. White and black arrowheads indicate the branch innervating the FDS. FDS, flexor digitorum superficialis; FDP, flexor digitorum profundus; MN, median nerve.

#### 3.2.2. Ulnar nerve

After coursing dorsal to the medial epicondyle of the humerus, the ulnar nerve gave off a muscular branch to the anconeus and then divided into two branches proximal to the forearm (Figure 3). One branch gave off muscular branches to the FDP and then communicated with a branch of the median nerve, as mentioned above. The other branch, regarded as the main trunk, ran between the FDP and the FCU and continued toward the palm, giving off muscular branches to the FCU. Tracing of the muscular branches from the median and ulnar nerves distally revealed that the ulnar nerve mainly supplied the ulnar and proximal part of the FDP, while the remaining part was innervated by the median nerve.

## 4. Discussion

In the present study, we examined the forearm flexors of koalas and identified distinctive morphological characteristics of the FDS, FDP, and their innervation patterns. The FDS was identified as a small muscle originating from the superficial surface of the FDP, and it divided into four thin distal tendons that extended to the digits II–V. In contrast, the FDP had a large muscle belly originating from the medial epicondyle, ulna, and radius, and its distal tendons extended to the digits I–V. These findings on muscle morphology were generally consistent with the descriptions of Young (1882) and Richards et al. (2023). Nevertheless, this study characterized and described the morphology of these muscles in more detail with fine photographs and schematic drawings, thereby more clearly demonstrating structural characteristics that had not been sufficiently illustrated in previous studies.

We further identified and described a distinctive innervation pattern in the median nerve, which gave off an ascending branch to the FDS. To our knowledge, this is the first report to describe this unique innervation pattern of the FDS among mammals. Previous studies were mostly limited to muscle morphology, and detailed descriptions of the innervation of these muscles were lacking. As mentioned in the Materials and Methods, considering muscle innervation patterns, in addition to muscle attachment sites and shape, is useful for interpreting muscle morphogenesis and homology (Anetai et al., 2026; Emura et al., 2020, 2023; Koizumi, 2019, 2022, 2023; Sakuraya *et al*., 2023; Velez-Garcia et al., 2026). Based on these findings, the present study discusses the morphology of the koala forearm flexors from the perspectives of comparative anatomy, morphogenesis, arboreal adaptation, and convergent evolution with primates.

### 4.1. Morphological and Functional Considerations of the FDS

Among primates, the FDS is generally a well-developed muscle that inserts into digits II–V and has a muscle belly as large as or larger than that of the FDP, although its proximal attachments vary among species (Emura et al., 2020, 2023; Hartman & Straus, 1969; Romer, 1997). For example, in marmosets and squirrel monkeys, the FDS originates only from the medial epicondyle of the humerus, whereas in spider monkeys it arises from both the medial epicondyle and the proximal ulna (Emura et al., 2020). The siamang FDS originates from the medial epicondyle and the central regions of the ulna and radius as in humans, whereas in western lowland gorillas and western chimpanzees it arises from the medial epicondyle and the central radius (Emura et al., 2023; Gray 1918).

In koalas, the FDS showed a particularly distinctive pattern of origin, lacking any proximal bony attachment and instead arising from the superficial surface of the FDP. Unlike in primates, the FDS had a much smaller muscle belly than that of the FDP. This reduced muscle belly size may be associated with the absence of a proximal bony attachment. A less-developed FDS relative to the FDP has also been reported in other marsupials. In several carnivorous marsupials, including quolls and dunnarts, the FDS is described as a tiny slip of muscle arising from the distal superficial aponeurosis of the FDP (Warburton and Marchal, 2017). Although the FDS in the southern brown bandicoot and bilby arises partly from the distal extremity of the humeral medial epicondyle, as well as from the fascia of the deep flexor mass, the muscle has nevertheless been described as having a small muscle belly that gives rise to very thin tendons (Warburton et al., 2013). In marsupial moles, the FDS is extremely reduced or nearly absent (Warburton, 2006). These findings suggest that the size and degree of development of the FDS vary among marsupial species, but that a relatively reduced FDS is not unusual among them. This morphology suggests that, unlike in primates, the koala FDS may function in close association with the FDP contraction rather than as an independent flexor. Thus, when the FDP contracts and its muscle belly becomes tense, the FDS is also likely to generate contractile force. In addition, the tendons inserting into the middle phalanges of digits II–V were extremely thin. Accordingly, the FDS alone is unlikely to exert a strong flexion force, and flexion of the proximal interphalangeal joints is considered to occur in an auxiliary or synchronized manner, accompanying the distal interphalangeal flexion produced by the FDP.

### 4.2. Morphological and Functional Considerations of the FDP

In koalas, the FDP arose from the medial epicondyle, ulna, and radius, gave rise to tendons extending to digits I–V, and had a much more clearly developed muscle belly than the FDS. This is consistent with the quantitative findings reported by Richards et al. (2023), in which the FDP had the largest normalized physiological cross-sectional area (PCSA), suggesting an architectural emphasis on force-generating capacity. These differences in the digital flexors may be attributable to the functional roles of claws during arboreal grasping. Many non-primate arboreal mammals grasp branches with hooked claws, whereas most primates, which have flat nails, grasp branches using prehensile hands and feet (Gebo, 2014; Kardong, 2012; Kent & Carr, 2001). Koalas, like other non-primate arboreal mammals, possess sharp claws at the tips of their digits. Because flexion of the distal interphalangeal joint is essential for hooking the claws onto branches, the FDP, which inserts into the distal phalanx, is likely to play a major functional role in this action. Its marked development in the koala may therefore reflect the substantial functional demand placed on this muscle during arboreal grasping.

In addition, no independent muscle corresponding to the FPL of great apes was observed in the koala. Therefore, the FDP component giving rise to the tendon inserting into digit I in koalas is considered to correspond to the FPL, suggesting that the FDP and FPL are present as a single muscle in the koala. It is known that in many non-human primates and non-primate mammals the FPL does not exist as an independent muscle (Hartman 1969; Honma and Sakai, 1992; Nickel et al., 1986). This suggests that the koala FDP retains the general mammalian pattern, although koalas have a unique digital arrangement: the space between digits II and III is widened, and digits I and II oppose the remaining digits (Figure 1).

### 4.3. Ascending branch of the median nerve to the FDS and its implications for morphogenesis

The koala FDS was innervated by branches originating from the main trunk of the median nerve near the carpal region and ascending proximally to the FDS. This unique extra-muscular course may reflect an underlying embryological process. A Previous study reported that, although the forelimbs begin to develop earlier, limb development is highly conserved across vertebrates and the observed changes are largely limited to shifts in timing and location (Keyte & Smith, 2010?). Accordingly, given the extensive developmental evidence from humans and experimental rodents, as well as the morphogenetic interpretations derived from anatomical studies, we interpret the koala FDS in the present study in light of the developmental processes of the FDS in these taxa.

In human fetuses, it has been reported that the proximal primordia of the FDS and palmaris longus arise in the forearm, whereas a distal primordium, namely the flexor digitorum brevis (FDB), appears near the palm (Gräfenberg, 1905). The definitive FDS is then thought to develop as this distal primordium migrates proximally and fuses with the proximal forearm primordia. Yamada (1986) also stated that the FDS comprises a ventral portion derived from the deep flexor musculature of the forearm together with additional muscle bundles that fuse within the palm. Given these findings, we infer that the koala FDS is derived predominantly from the FDB. This is because the koala FDS has no bony origin from the medial epicondyle of the humerus, radius, or ulna. Accordingly, it is unlikely to be derived from the proximal forearm primordia of the deep flexor compartment. The unique ascending course of the muscular branch from the median nerve to the koala FDS could be consistent with a developmental scenario in which this branch entered the muscle when the distal primordium was located near the carpal region, after which the muscle ascended together with its supplying branch. This unusual course of the median nerve and the morphology of the muscular branch supplying the FDS may represent distinctive features in koalas. To our knowledge, no such ascending course of the muscular branch to the FDS has been reported in the available comparative anatomical literature.

The absence of fusion between the koala FDS and the proximal forearm primordia, if present, may be related to differences in forelimb development and function between primates and marsupials. As noted above, although the developmental process of the marsupial forelimb is generally comparable to that of other vertebrates, differences in the relative timing and spatial organization of limb development may have influenced the formation of the FDS. Species-specific developmental events during early marsupial forelimb development may also have contributed to the FDS formation. Although the details of the developmental mechanisms underlying this morphology and its phylogenetic background remain unclear, the present findings may provide useful insights into the neuroembryology and comparative embryology of the forearm flexors.

## Conclusion

In this study, we identified the morphological characteristics and distinctive innervation pattern of the forelimb digital flexors in koalas and discussed them in comparison with those of primates adapted to arboreal life. The results suggest that the forelimb digital flexors, particularly the FDS, differ from those of primates not only in morphology but also in morphogenetic background. Thus, koalas and primates appear to have achieved adaptation to arboreal life through different strategies within the forearm musculature.

## Acknowledgement

We would like to express our deepest gratitude to Prof. Masahiro Koizumi (Tokyo Ariake Medical University) for providing koala specimens and allowing to conduct dissections.

## Author contributions

**DS** data curation, writing–original draft, writing–review and editing. **SA** conceptualization, methodology, data curation, validation, writing–original draft, writing–review and editing, supervision. **HA** validation, writing–review and editing. **JK** and **CL** resource, writing-review and editing. **KT** supervision, writing–review and editing. **ST** resource, writing-review and editing, project administration.

